# CHD4 slides nucleosomes by decoupling entry- and exit-side DNA translocation

**DOI:** 10.1101/684860

**Authors:** Yichen Zhong, Bishnu Prasad Paudel, Daniel P. Ryan, Jason K. K. Low, Charlotte Franck, Karishma Patel, Max J. Bedward, Richard J. Payne, Antoine M. van Oijen, Joel P. Mackay

## Abstract

Chromatin remodellers hydrolyse ATP to move nucleosomal DNA against histone octamers. The mechanism, however, is only partially resolved, and unclear if it is conserved among the four remodeller families. Here we use single-molecule assays to examine the mechanism of action of CHD4, which is part of the least well understood family of remodellers. We demonstrate that the binding energy for CHD4-nucleosome complex formation – even in the absence of nucleotide – triggers significant conformational changes in DNA at the entry side, effectively priming the system for remodelling. During remodelling, flanking DNA enters the nucleosome in a continuous, gradual manner but exits in concerted 4–6 base-pair steps. This decoupling of entry- and exit-side translocation suggests that ATP-driven movement of entry-side DNA builds up strain inside the nucleosome that is subsequently released at the exit side by DNA expulsion. We propose a mechanism for nucleosome sliding based on these and published data.

## Introduction

The nucleosome is the fundamental unit of the eukaryotic genome. Each nucleosome is made up of a histone octamer core containing two copies of each histone (H2A, H2B, H3 and H4) and wrapped by 147 base pairs (bp) of DNA (Kornberg, 1974). The histone-DNA interface consists mainly of inward-facing sections of the DNA minor groove, which are defined as superhelical locations (SHL) ± 0.5–6.5; the adjacent outward-facing minor grooves are named SHL ± 1–6 (Flaus, 2011) (Figure 1A). The nucleosome provides a folding scaffold that allows metres of linear DNA to be packed inside micron-sized nuclei. Through the activity of chromatin modifying enzymes, nucleosomes also regulate DNA accessibility and therefore gene expression.

**Figure 1.**
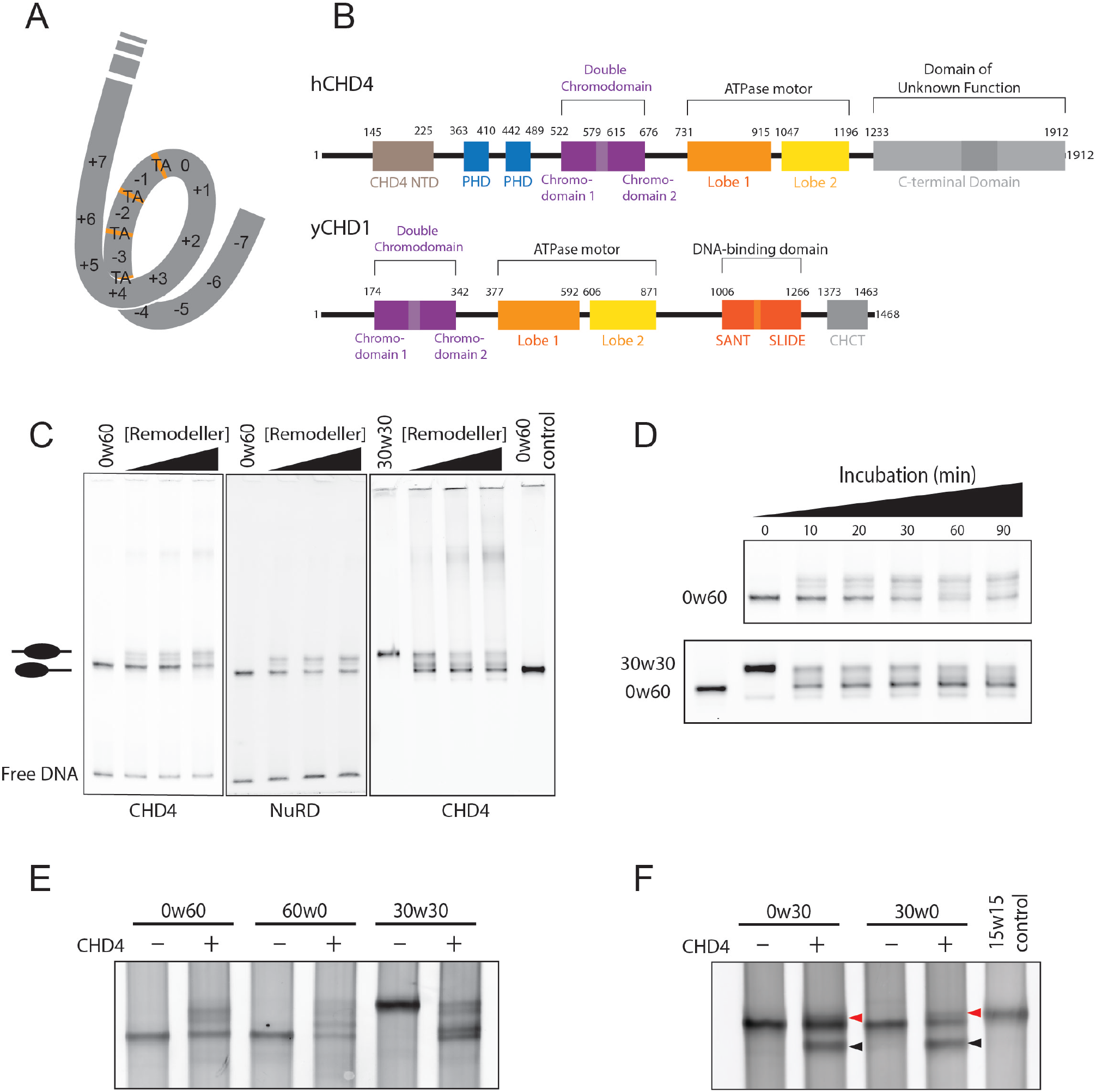
CHD1/CHD4 domain topology and nucleosome repositioning assays. **A.** Schematic showing the fifteen SHL sites in the 601 nucleosome positioning sequence. Orange bands indicate four phased TpA dinucleotides that are spaced 10 bp apart. **B.** Domain architectures of yeast CHD1 and human CHD4 with residues at domain boundaries indicated (NTD: <u>N</u>-terminal <u>d</u>omain, PHD: <u>P</u>lant <u>h</u>omeo<u>d</u>omain, CHCT: <u>CH</u>D1 <u>C</u>-terminal domain). **C.** Gel-based nucleosome repositioning assays carried out with the indicated nucleosomes and remodellers. Fluorescently labelled nucleosomes were treated with the indicated remodeller for 60 min, the reaction was stopped by adding dsDNA (33 µg/mL) and then the samples were run on 5% native polyacrylamide gels. Symmetrically positioned nucleosomes are retarded relative to asymmetrically positioned species, as indicated. **D.** Gel-based nucleosome repositioning assays carried out as described in (C), except that an incubation time-course was carried out at a single CHD4 concentration (5 nM). Assays using 0w60 (*upper panel*) or 30w30 (*lower panel*) substrates are shown; a 0w60 control lane was included in the lower panel. **E** and **F.** Nucleosome sliding assays carried out as described in (C), using the indicated nucleosome substrates and 5 nM CHD4. Remodelled products (0w30 and 30w0) are indicated by red arrows, and the possible hexosome band is indicated by black arrows.

Chromatin remodellers are specialized translocases that use the energy of ATP hydrolysis to reposition, eject, and replace histones within the nucleosome. These events happen in response to stimuli such as epigenetic modifications and cell cycle signals (Narlikar et al., 2013), thus exposing the DNA to (or protecting it from) other DNA-binding or transcription regulatory factors (Clapier and Cairns, 2009). Mutation or dysfunction of chromatin remodellers often leads to severe consequences, including disrupted cell cycle, tumorigenesis and even early embryonic lethality (Langst and Manelyte, 2015).

All known chromatin remodellers are superfamily 2 (SF2) ATPase motors, which consist of two RecA-like lobes that form an active-site cleft (Flaus et al., 2006, Cairns, 2007). However, the domains flanking this conserved ATPase domain vary and are used to classify chromatin remodellers into four families: SWI/SNF (mating type <u>swi</u>tching/<u>s</u>ucrose <u>n</u>on-<u>f</u>ermenting), ISWI (<u>i</u>mitation <u>swi</u>tch), INO80 (<u>ino</u>sitol), and CHD (<u>c</u>hromodomain <u>h</u>elicase <u>D</u>NA-binding). Unique among these families, CHD remodellers possess two tandem chromodomains adjacent to the ATPase. Some CHD remodellers can function as a monomer, such as yeast CHD1 (Ramirez et al., 2012), but metazoan CHDs generally exist in multi-protein complexes. The CHD family member CHD4 is a subunit of the nucleosome remodelling and deacetylase (NuRD) complex (Torchy et al., 2015); together, CHD4 and the histone deacetylases HDAC1 and −2 provide NuRD with both nucleosome remodelling and histone deacetylation activities.

The two chromodomains in CHD1 and CHD4 regulate the ATPase activity (Hauk et al., 2010, Watson et al., 2012) and are essential for recognising methylated lysine in histone H3 or associating with chromatin (Flanagan et al., 2005, Yap and Zhou, 2011, Ramirez et al., 2012). Further, the N-terminal portion of CHD4 contains two N-terminal plant homeodomains (PHD) that can interact with trimethylated H3K9 and unmodified H3K4 (Mansfield et al., 2011). At the C-terminal end, the DNA-binding SANT-SLIDE domain (DBD) in CHD1, which unpeels ~30 bp of DNA from the histone octamer surface (Ryan et al., 2011), is replaced by a large uncharacterized C-terminal domain (CTD) in CHD4 (Figure 1B). Both the PHDs and the CTD are essential for the repression activity of the NuRD complex (Mansfield et al., 2011, Ramirez et al., 2012). The exact significance of having both chromodomains and PHDs for H3 recognition is unknown, but this structural feature is also commonly seen in other chromatin remodelling proteins and might be important in connecting the recognition of histone modifications and the chromatin remodelling activity (Mansfield et al., 2011).

Even in the absence of other NuRD subunits, CHD4 can reposition an asymmetric mononucleosome substrate bearing different lengths of DNA at the two sides of the histone octamer, yielding a more symmetric structure (Levendosky et al., 2016, Kovac et al., 2018). However, this activity is much weaker in comparison to CHD1 (Watson et al., 2012). To date, the mechanism of this ATPase-driven DNA movement remains ambiguous, but two popular models have been proposed (Saha et al., 2006, Bowman, 2010). The “DNA loop/wave propagation” model states that the DNA entering the nucleosome from the entry side forms a ~10-bp bulge/loop on the octamer surface that quickly propagates around the histone core and is subsequently released from the exit site. The “twist diffusion” model proposes that the remodeller changes the DNA helix structure and length, causing a twist defect, and hydrolysis of ATP results in directional transfer of a few base pairs to the adjacent DNA segment (Winger et al., 2018, van Holde and Yager, 2003).

In this study, we investigated the mechanism by which CHD4 remodels mononucleosomes. We have combined bulk gel-based assays and single-molecule Förster resonance energy transfer (smFRET) methods to monitor the nucleosome sliding activity of CHD4 in real time. The data show that during remodelling, DNA enters the nucleosome in a continuous manner but exits in bursts of 4−6-bp, demonstrating that the entry and exit processes are partially decoupled. We also show that CHD4 binding in the absence of nucleotide induces substantial dynamics at the entry side of the nucleosome, suggesting that the binding energy of CHD4 alone makes a significant contribution to the remodelling process. Overall, our results reveal mechanistic aspects of the process by which CHD4 re-organizes nucleosomes in the genome.

## Results

### CHD4 remodels both symmetric and asymmetric nucleosome substrates

We first assessed the nucleosome sliding activity of human CHD4 using a gel-based remodelling assay. We expressed and purified human histones H2A, H2B, H3 and H4 and reconstituted nucleosomes using the 601 Widom positioning DNA sequence (Lowary and Widom, 1998) to which a 60-bp extension was added at the +7 end (Figure 1A). We refer to this construct as 0w60. Following incubation with CHD4 and ATP, the reaction mixture was run on a native gel. New bands were observed that represent more symmetrically positioned nucleosomes, which have been shown previously to migrate more slowly (Hamiche et al., 1999, Langst et al., 1999) (Figure 1C, *left panel*). Although the intensity of the upper bands increased at higher CHD4 concentrations and with longer incubation times (Figure 1D, *top panel*), we still observed a considerable proportion (~50%) of starting material. We repeated the remodelling assay using the same amount of native NuRD complex, purified as previously described (Low et al., 2016), which in addition to CHD4 contains HDAC, MTA, RBBP, MBD and GATAD2 subunits. An identical pattern of shifted bands was observed (Figure 1C, *middle panel*), indicating that the additional subunits do not significantly alter the distribution of products, in contrast to the observation made previously for the CHRAC complex (Eberharter et al., 2001).

To investigate the effect of substrate conformation, and to determine if CHD4 could reversibly remodel its products, we also assessed the activity of CHD4 towards a nucleosome bearing 30-bp extensions on both sides of the nucleosome core particle (30w30). In this case, ~80% of the starting material was converted to less symmetric products (Figure 1C, *right panel*, and Figure 1D, *lower panel*), indicating that CHD4 can remodel mononucleosomes with either symmetric or asymmetric extensions, but displays a substrate preference and greater remodelling efficiency towards nucleosomes with extensions on both sides.

The 601 Widom sequence used for our nucleosomes is non-palindromic, as its two SHL2 sites have significant sequence differences. The SHL–2 site (Figure 1A) contains four phased TpA dinucleotides at inward-facing minor groove positions, providing more DNA-nucleosome stability (Chua et al., 2012, Ngo et al., 2015). Therefore, we reconstituted a 60w0 construct so that the histone octamer would be remodelled towards the other end of the 601 sequence. Like its 0w60 counterpart, 60w0 was partially but not fully remodelled. With the extension now located on the ‘minus’ side, closest in sequence to the TA-rich region, the final set of products displayed a similar but not identical distribution (Figure 1E). The bands with intermediate migration probably represent partially remodelled nucleosomes. These data suggest that CHD4 has some ability to discriminate between substrates with different DNA sequences.

Next, we tested the effect of the extra-nucleosomal DNA length on CHD4 remodelling activity. Remodellers such as INO80 and ISWI can sense the length of extra-nucleosomal DNA and have a preference for substrates with longer flanking DNA (Zhou et al., 2018, Kagalwala et al., 2004). CHD4 was able to remodel nucleosomes with a 30-bp extension (0w30 and 30w0) (Figure 1F), although the extent of remodelling was significantly less than for the 0w60 substrate. A lower-migrating band also appeared after incubation and, based on published data (Levendosky et al., 2016), is likely to be a hexasome that arises from loss of a single H2A-H2B dimer. We are unsure whether the hexasome is a by-product from remodelling of an asymmetric nucleosome with shorter flanking DNA or rather represents a previously undescribed activity of CHD4. A 30w30 substrate showed significantly more remodelling and less hexasome formation (Figure 1E), suggesting that total DNA length and/or sequence might impact on remodelling behaviour and also the stability of nucleosome during remodelling.

### The CHD4-nucleosome interaction is independent of flanking DNA length

Because our remodelling data suggested that a symmetric nucleosome with longer flanking DNA is a more favourable substrate for CHD4, we sought to test whether this preference arises from DNA-binding preferences of the protein. We therefore bound FLAG-CHD4 to anti-FLAG beads and incubated the beads with 0w0, 0w60, 30w30 or 60w60 nucleosomes. Surprisingly, all constructs were pulled down by CHD4 with no clear preference for the flanking DNA lengths and/or substrate symmetry (Figure 2A), suggesting the differences in remodelling activity are not driven by substrate binding selectivity.

**Figure 2.**
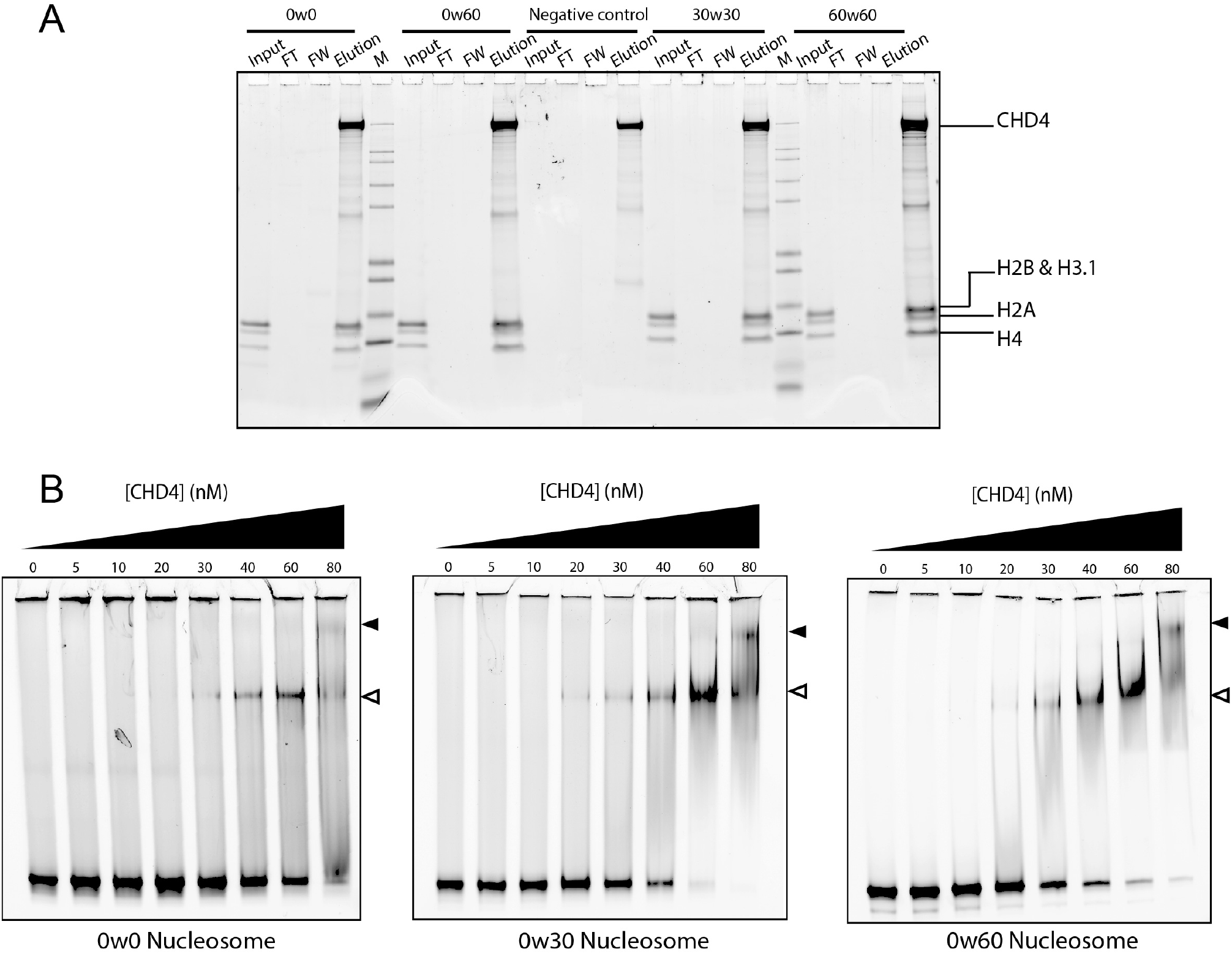
The affinity of CHD4 for DNA is not strongly dependent on the presence of flanking extra-nucleosomal DNA. **A.** Pulldowns in which FLAG-CHD4 is immobilized on FLAG-Sepharose beads and incubated with the indicated nucleosomes, followed by elution with FLAG peptide. Nucleosome input, unbound nucleosome/flow through (FT), final wash (FW) and elutions of each nucleosome construct were analysed by SDS-PAGE, along with a size indicator, Mark12 Standard (M). The negative control contains no nucleosome in the input. **B.** EMSAs showing the binding of CHD4 to the indicated nucleosomes. Nucleosome concentration was 90 nM (0w0) or 60 nM (0w30 and 0w60). Open and filled triangles indicate complexes that probably contain one or two CHD4 molecules, respectively.

Electrophoretic mobility shift assays (EMSAs) were then used to quantify DNA-binding affinity. Figure 2B shows that 0w0, 0w30 and 0w60 all bind CHD4 with comparable affinity (with dissociation constants of ~40–80 nM. The fact that even the nucleosome with no flanking DNA (0w0) displayed a similar affinity to the other constructs suggests that a significant number of the contacts made by CHD4 are to the nucleosome core particle rather than to flanking DNA. Furthermore, binding of a second CHD4 molecule can be observed at high CHD4 concentration, indicating that two CHD4 binding sites exist and that these two sites must lie sufficiently far from the dyad axis of the nucleosome that they can be co-occupied. The pattern of shifts also indicates that the two CHD4 binding events are not cooperative.

### CHD4 ejects DNA from the nucleosome in multi-base-pair steps

To further understand the mechanism of CHD4-driven nucleosome sliding, we used single-molecule FRET (smFRET) to monitor nucleosome conformational changes. The ability to follow a single nucleosome through time allows the observation of transient intermediates during the remodelling process and thus provides access to mechanistic details of the sliding process. We assembled nucleosomes using DNA tagged with a 5’ donor dye (AlexaFluor555, AF555) at one end and with 5’ biotin at the other (Figure 3A). We used the well-studied T120C mutant of H2A in these experiments, which was coupled to the acceptor dye AlexaFluor647 (AF647) via the cysteine. We optimised the labelling reaction to be incomplete (50% of H2A labelled) so that a large portion of nucleosomes contained only one copy of labelled H2A at positions that are either proximal or distal to the AF555 DNA tag. We named these constructs *n*^AF555^w60^Bio^, where *n* indicates the number of base pairs between the AF555 tag and the 601 sequence. Because our bulk gel-based experiments suggested that CHD4 remodels 0w60 to a more symmetric conformation, we refer the end with the 60-bp flanking DNA as the entry site, and the other as the DNA exit site. The nucleosome substrates were immobilized on a streptavidin-PEG-coated coverslip (Figure 3A) embedded in a microfluidic channel, and the coverslip was imaged using total internal reflection fluorescence (TIRF) microscopy. The AF555 and AF647 dyes form a FRET pair, which reports on the distance between the DNA and histone label positions.

**Figure 3.**
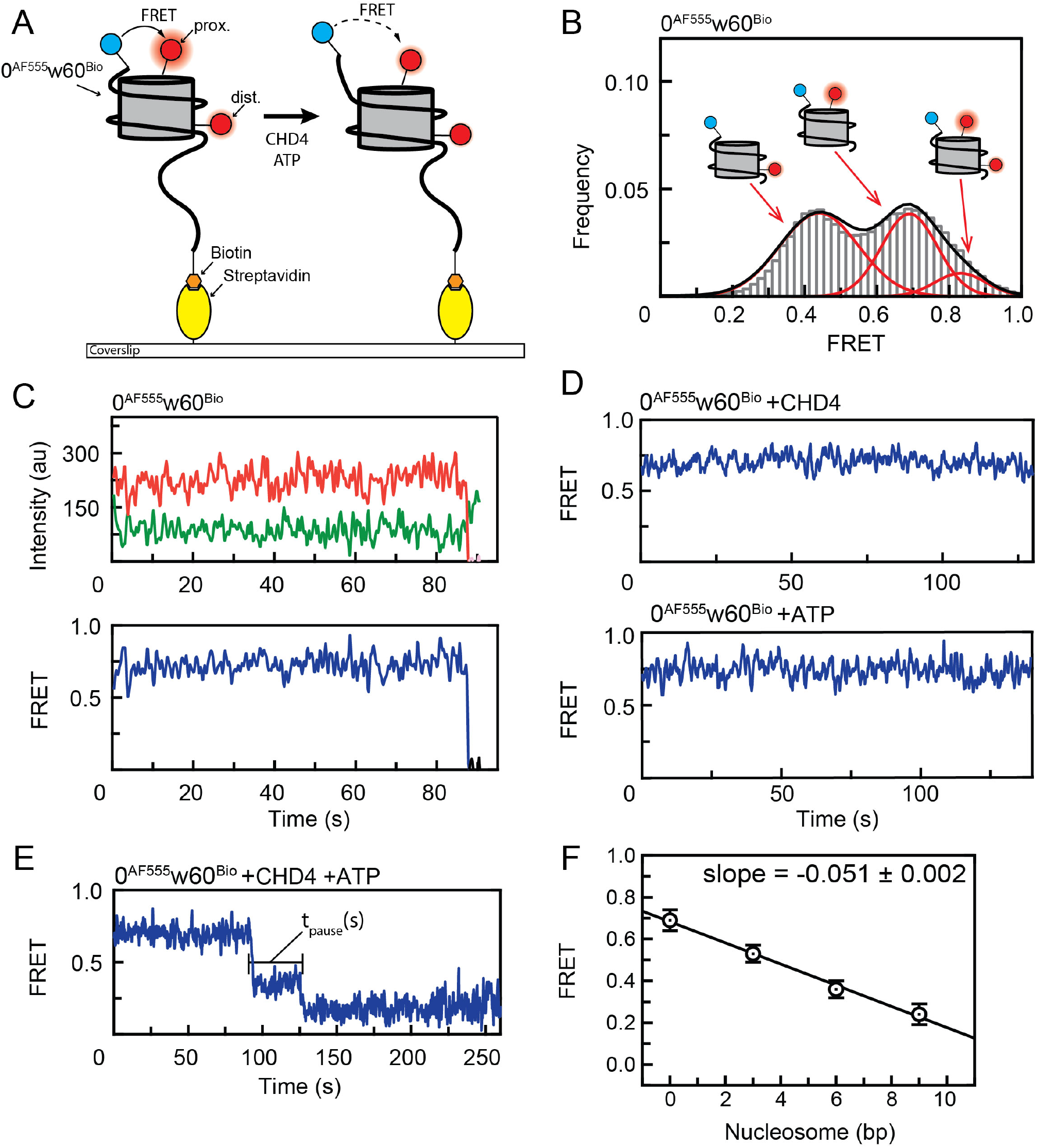
A single-molecule FRET (smFRET) assay for nucleosome sliding shows that CHD4 drives multi-base-pair movements of nucleosomal DNA at the exit side. **A.** Diagram depicting the setup for the smFRET assay and nucleosome substrate before and after remodelling by CHD4. A 0^AF555^w60^Bio^ nucleosome containing either proximal or distal or both AF647-labelled H2A is assembled onto a PEGylated coverslip via biotin at the longer end of the flanking DNA (AF555 and AF647 are represented by red and blue circles respectively). FRET is then monitored as a function of time under different conditions. **B.** Pre-reaction distribution of nucleosomal FRET states for 0^AF555^w60^Bio^ nucleosomes. Low, mid- and high FRET states correspond to particles containing H2A in the distal, proximal or both positions (relative to the DNA-bound AF555), respectively. **C.** FRET vs time trace of 0^AF555^w60^Bio^ nucleosome bearing a proximal H2A label. Donor AF555 fluorescence (green), acceptor AF647 fluorescence (red) and FRET (blue) were shown. **D.** FRET vs time traces of 0^AF555^w60^Bio^ nucleosome bearing a proximal H2A label in the presence of 2 nM CHD4 (*top*) or 1 mM ATP (*bottom*). **E**. Time dependence of FRET for a single 0^AF555^w60^Bio^ particle following the addition of 2 nM CHD4 and 200 µM ATP. Two clear drops in FRET are observed. We define the pause time *t*_*pause*_ as the duration between two FRET changes. **F.** Calibration of FRET values for *n*^AF555^w60^Bio^ nucleosomes. The FRET of proximally labelled particles was measured as a function of the number of base pairs (*n*) added at the DNA exit site and mid-FRET peak values for each construct were obtained by fitting to a Gaussian distribution. Plotting the change in mid-FRET value as a function of *n* yielded a slope of –0.051± 0.002. Error bars represent standard deviation of the fit from at least two independent measurements.

Imaging of immobilized 0^AF555^w60^Bio^ nucleosomes yielded three populations with distinct FRET values, which correspond to nucleosomes bearing AF647-H2A at either the proximal or the distal site, relative to AF555, or both (Figure 3B). Unless indicated, only the mid-FRET population that contains a single, proximally located label was selected. To establish the behaviour of this system as a function of time, we first monitored the FRET of the immobilized 0^AF555^w60^Bio^ nucleosomes alone. Figure 3C shows that no significant change in FRET is observed until one of the fluorophores undergoes photobleaching. Similarly, introducing CHD4 or ATP alone does not give rise to any changes (Figure 3D), indicating that the binding of CHD4 or the presence of ATP does not induce any significant conformational changes in the nucleosome that alter the distance between the two fluorophores.

Next, we treated 0^AF555^w60^Bio^ nucleosomes with both CHD4 and ATP in a continuous flow manner and monitored FRET as a function of time. A stepwise reduction in FRET was seen, indicating that the two fluorophores had moved away from each other (Figure 3E and **Supplementary Figure 1A**). The traces consistently showed two distinct and sharp drops in FRET, and these two drops were separated by a period of time (the pause time, *t*_*pause*_) during which FRET remains constant. In some cases, transient excursions to lower FRET values for up to ~10 s were observed (changes of up to ~0.5 FRET units; see for example **Supplementary Figure 1B**), followed by a return to the pre-excursion value. A similar overall pattern of two drops in FRET was observed for particles containing a distally labelled H2A (**Supplementary Figure 1C**).

To correlate the observed changes in FRET to translocation of a particular number of base pairs of DNA, we reconstituted a series of *n*^AF555^w60^Bio^ nucleosomes with *n* = 0, 3, 6 or 9 and constructed a calibration curve (Figure 3F). These data indicated that a reduction in FRET of 0.051±0.002 units corresponds to a 1-bp change in AF555 position relative to the H2A-bound AF647. The two drops in Figure 3E therefore correspond to 6- and 4-bp translocations (for the first and second steps, respectively) of the labelled end of the DNA away from the core particle. As the DNA helix is around 10 bp per turn, these step sizes suggested that roughly a half-turn of DNA moves out of the nucleosome at each step and that, after two steps, a full turn of DNA will be translocated.

### The pause time t_pause_ is ATP concentration dependent

To probe the mechanism of remodelling, we next assessed the dependence of the pause time between the two remodelling events (*t*_*pause*_, Figure 3E) on reaction conditions. We first varied the CHD4 concentrations but found that a 100-fold change in concentration (from 200 pM to 20 nM) resulted in only a small increase in *t*_*pause*_ (from a mean of 0.7 to 0.8 s, Figure 4A), suggesting that the remodelling process does not rely on the dissociation and rebinding of CHD4.

**Figure 4.**
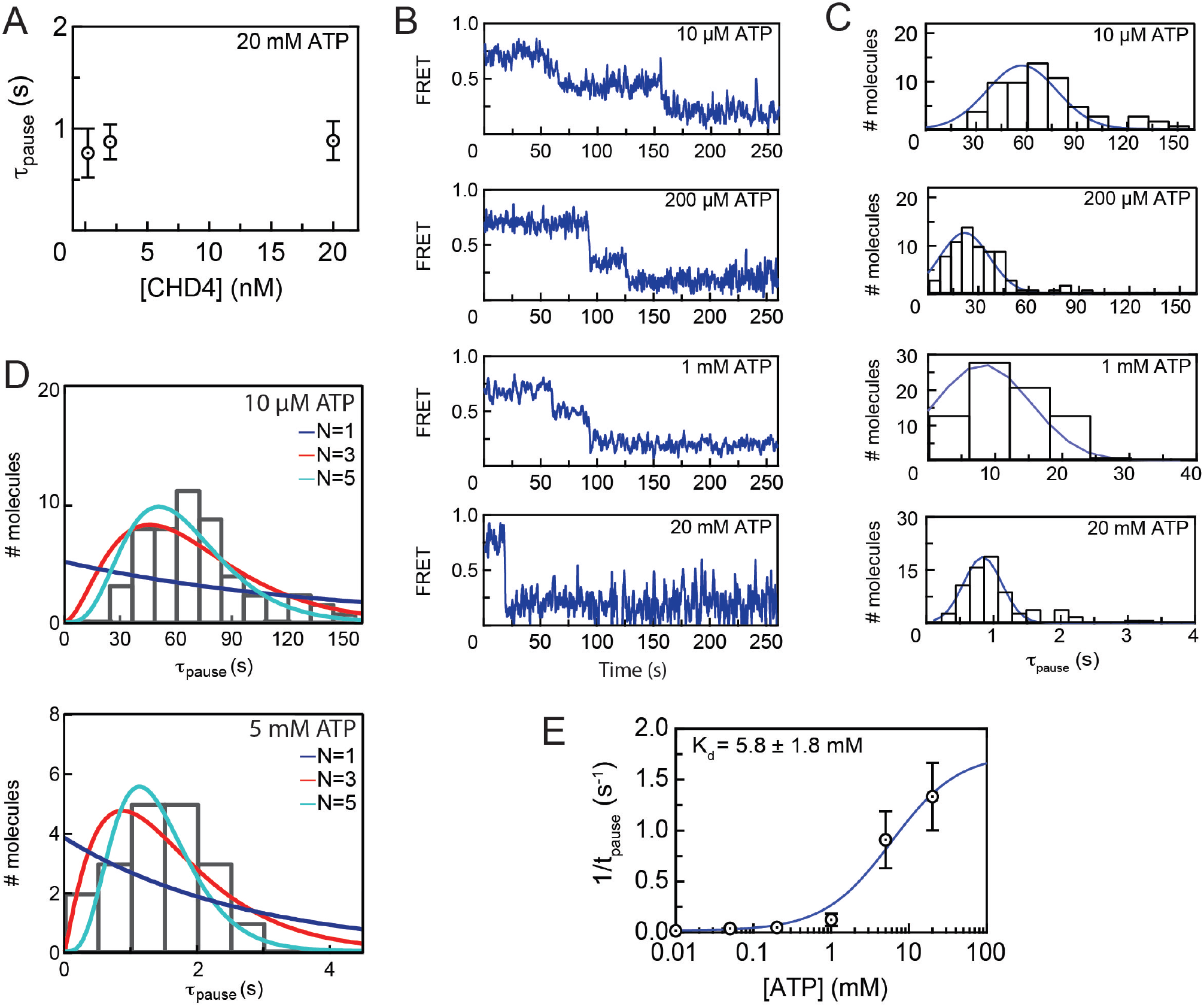
CHD4-mediated nucleosome sliding is processive and depends on the binding of multiple ATP molecules. **A.** Distribution of *t*_*pause*_ times for remodelling of 0^AF555^w60^Bio^ at the indicated CHD4 concentrations and 20 mM ATP. Error bars represent standard deviation (of 45, 74 and 56 molecules at 0.2, 2 and 20 nM CHD4, respectively). **B.** Typical traces from smFRET assays carried out with 2 nM CHD4 and at the indicated ATP concentrations. **C.** Histograms showing *t*_*pause*_ distributions of 70–82 particles from the experiments shown in (B). The histograms are fitted to a Gaussian distribution. **D**. Pause time histograms for experiments carried out at 10 μM ATP (*top*) or 5 mM ATP (*bottom*) are overlaid with gamma distributions depicting different numbers of fundamental reaction steps (N = 1–3). **E.** A 1:1 binding isotherm fit of the mean *t*_*pause*_ time as a function of ATP concentration, with data taken from the assays in (C). Error bars represent standard deviation.

In contrast, the reaction was strongly dependent on ATP concentration. Increasing ATP concentration from 10 µM to 20 mM reduced *t*_*pause*_ by roughly 60-fold, to the point where the separation between two FRET drops was almost unobservable (Figure 4B-C). Thus, under these conditions, ATP binding (and probably hydrolysis) is the rate-limiting step of the reaction. Importantly, if the transition between the two FRET drops comprises a single fundamental reaction step, then the distribution of *t*_*pause*_ values in each reaction should follow an exponential function. If the pause period instead comprises a number of sequential reaction steps, *t*_*pause*_ should follow a gamma function (Floyd et al., 2010). Figure 4C shows that the distribution of *t*_*pause*_ is not exponential at any of the ATP concentrations tested. Fitting the *t*_*pause*_ distribution at 10 μM and 5 mM ATP to a gamma function indicates that this part of the remodelling reaction involves approximately 5 reaction steps (Figure 4D).

Given the dependence of *t*_*pause*_ on ATP concentration, we conclude that the pause period comprises ~5 ATP binding and hydrolysis events, which is approximately equivalent to the number of base pairs being translocated at the exit site. We also plotted 1/*t*_*pause*_ against ATP concentration and fitted a simple 1:1 binding isotherm (Figure 4E); the fit provides a pseudo-affinity of ATP for CHD4 of 5.8 ± 2 mM, which is within the range of estimated intracellular ATP concentration of 1–10 mM (Beis and Newsholme, 1975). Together, these data show that CHD4 is a processive remodeller that hydrolyses at least 5 ATP molecules per translocation step at the exit side.

### CHD4 binding introduces significant dynamics at the DNA entry side, which are further enhanced in the presence of ATP

Up to this point, we had only examined the behaviour of the DNA at the exit side of the 0^AF555^w60^Bio^ nucleosome. To probe the behaviour of the entry side, we constructed nucleosomes with the AF555 donor located 9 bp into the 60-bp extension on the entry side [0w(9^AF555^)60^Bio^ nucleosomes, Figure 5A]. In this case, most particles displayed FRET values of 0.2 or 0.4 (**Supplementary Figure 2A**), reflecting the increase in distance between the two fluorophores. The two values most likely correspond to the presence of a distal and proximal AF647-labelled H2A (relative to the position of the AF555 label on the entry side DNA), respectively. Because we were expecting the FRET of 0w(9^AF555^)60^Bio^ to increase during CHD4 remodelling, we used the distally labelled particles for our remodelling analysis, in order to maximize the resolution available.

**Figure 5.**
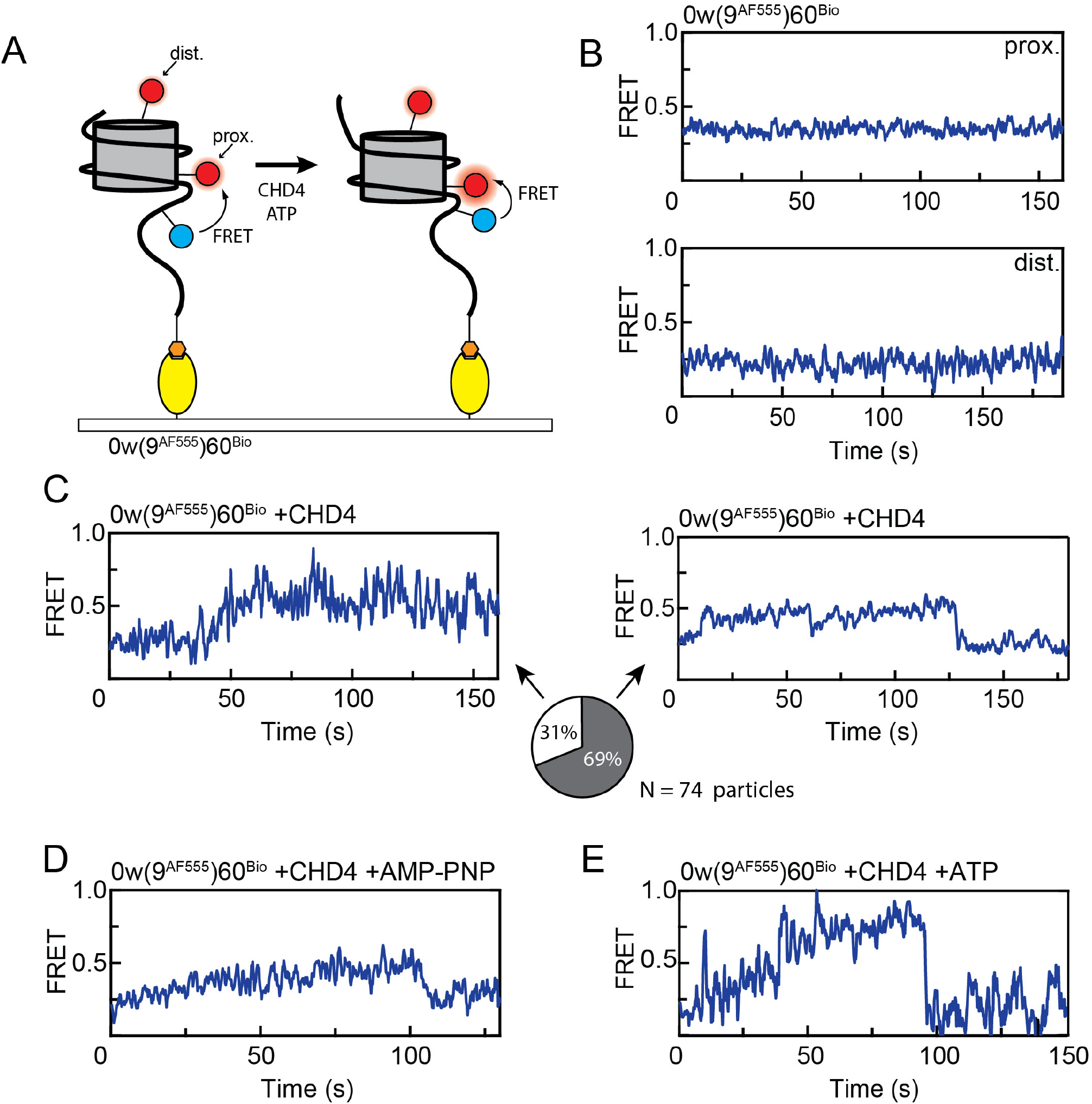
CHD4 binding induces changes in extra-nucleosomal DNA at the TA-poor side. **A.** Schematic showing the labelling scheme for 0w(9^AF555^)60^Bio^ and nucleosomal conformations before and after CHD4 remodelling. **B.** FRET vs time traces of 0w(9^AF555^)60^Bio^ alone with a proximal (*top*) and distal (*bottom*) AF647-labelled H2A. **C.** FRET vs time trace of 0w(9^AF555^)60^Bio^ bearing a distal AF647 label, showing a gradual increase upon the addition of 2 nM CHD4. This newly established structure can be relatively stable (*left*) or can be transient, dropping back to the initial state (*right*). The proportions of each scenario (from 74 molecules) are illustrated as a pie chart. **D.** FRET vs time trace of 0w(9^AF555^)60^Bio^ bearing a distal AF647 label, showing an increase upon the addition of 2 nM CHD4 and 1 mM AMP-PNP. **E**. FRET vs time traces of 0w(9^AF555^)60^Bio^ bearing a distal AF647 label showed a greater increase during CHD4 remodelling in presence of ATP, comparing to the changes induced by CHD4 alone.

In line with our previous observations for 0^AF555^w60^Bio^, the 0w(9^AF555^)60^Bio^ nucleosome alone did not undergo any significant FRET changes as a function of time (Figure 5B). However, in sharp contrast, the addition of CHD4 in the absence of ATP resulted in significant time-dependent changes in FRET. A gradual increase in FRET gave rise over the course of ~5–50 s to a long-lived state with a FRET value of ~0.45 (Figure 5C). Interestingly, for the majority of the molecules (69%, 51 out of 74 particles), the FRET subsequently returned to its starting value, via what was relatively a faster transition than the initial rise, whereas many of the remaining particles exhibited the higher FRET state until fluorophore bleaching occurred (Figure 5C). Similar behaviour was observed in the presence of the ATP analogue AMP-PNP (Figure 5D).

To observe entry-side behaviour during remodelling, we treated the 0w(9^AF555^)60^Bio^ nucleosomes with 2 nM CHD4 and 1 mM ATP. Under these conditions, nucleosomes containing a distal H2A label consistently displayed a gradual increase in FRET from 0.2 to ~0.6–0.8, a significantly greater change than that observed with CHD4 alone. Of the nucleosomes that underwent this increase, 64% subsequently exhibited a sudden return to ~0.2 (Figure 5E). In some cases, this cycle was repeated over the timescale of our observation. The increase in FRET is consistent with a gradual movement of the entry-side DNA into the nucleosome core in the presence of ATP, which would shorten the distance between the fluorophores. Nucleosomes bearing a proximally labelled H2A behaved similarly, with their FRET states increased from 0.4 to ~0.6–0.7 upon CHD4 binding (**Supplementary Figure 2B**) and a greater increase to ~0.8 in presence of CHD4 and ATP (**Supplementary Figure 2C**), both followed by a return to ~0.4.

We interpret the sharp drop in FRET observed for these nucleosomes as ‘reversions’ in which the DNA returns to its starting extra-nucleosomal position due to unsuccessful remodelling. Gel-based remodelling assays show that this entry-side labelled nucleosome could not be remodelled to the same extent as unlabelled or exit-side labelled nucleosomes (**Supplementary Figure 3**), perhaps due to steric hindrance caused by the bulky fluorophore on the DNA.

These data show that the binding of CHD4 – either alone or in the presence of the nucleotide analogue AMP-PNP – induces dynamic changes in nucleosome structure at the entry side and that ATP-driven remodelling proceeds with a gradual increase in FRET at the entry side alongside discrete downward jumps at the exit side.

### The effect of CHD4 binding on nucleosome dynamics is observed on both sides of the nucleosome

The observations above suggest that binding of CHD4 causes a change in the conformation of the extra-nucleosomal DNA on the entry side (or TA-poor side; Figure 1B) – a change that decreases the distance between the two fluorophores. One possible explanation is that DNA at the entry side translocates a short distance into the nucleosome even upon remodeller binding in the absence of nucleotide. A similar effect has been observed for CHD1; remodeller binding caused a 1–3 bp shift of the DNA in the TA-poor region (Winger et al., 2018).

Because our gel-based data indicate that the nucleosome has two CHD4 binding sites and that CHD4 can remodel both 0w60 and 60w0 substrates, we hypothesize that remodelling can occur in either direction, and that the direction depends on the site of CHD4 binding. Thus, either end of the nucleosome can act as an entry side providing it has sufficient extra-nucleosomal DNA. To test this idea, we assembled nucleosomes using a 9^AF555^w60^Bio^ DNA sequence (Figure 6A). Again, the nucleosome alone displayed no significant dynamic behaviour (Figure 6B) but the addition of CHD4 caused an increase in FRET (for 78 out of 91 molecules) from 0.2 to ~0.4 for distally labelled nucleosomes; proximal labelled nucleosomes increased from 0.4 to ~0.6 (Figure 6C). Out of these 78 molecules, 34 returned to the initial FRET state during the timescale of our measurement, whereas the remaining 44 remain at the higher FRET value. These data show that CHD4 elicits similar changes to flanking DNA at both the TA-rich and TA-poor ends of the 601-based nucleosome.

**Figure 6.**
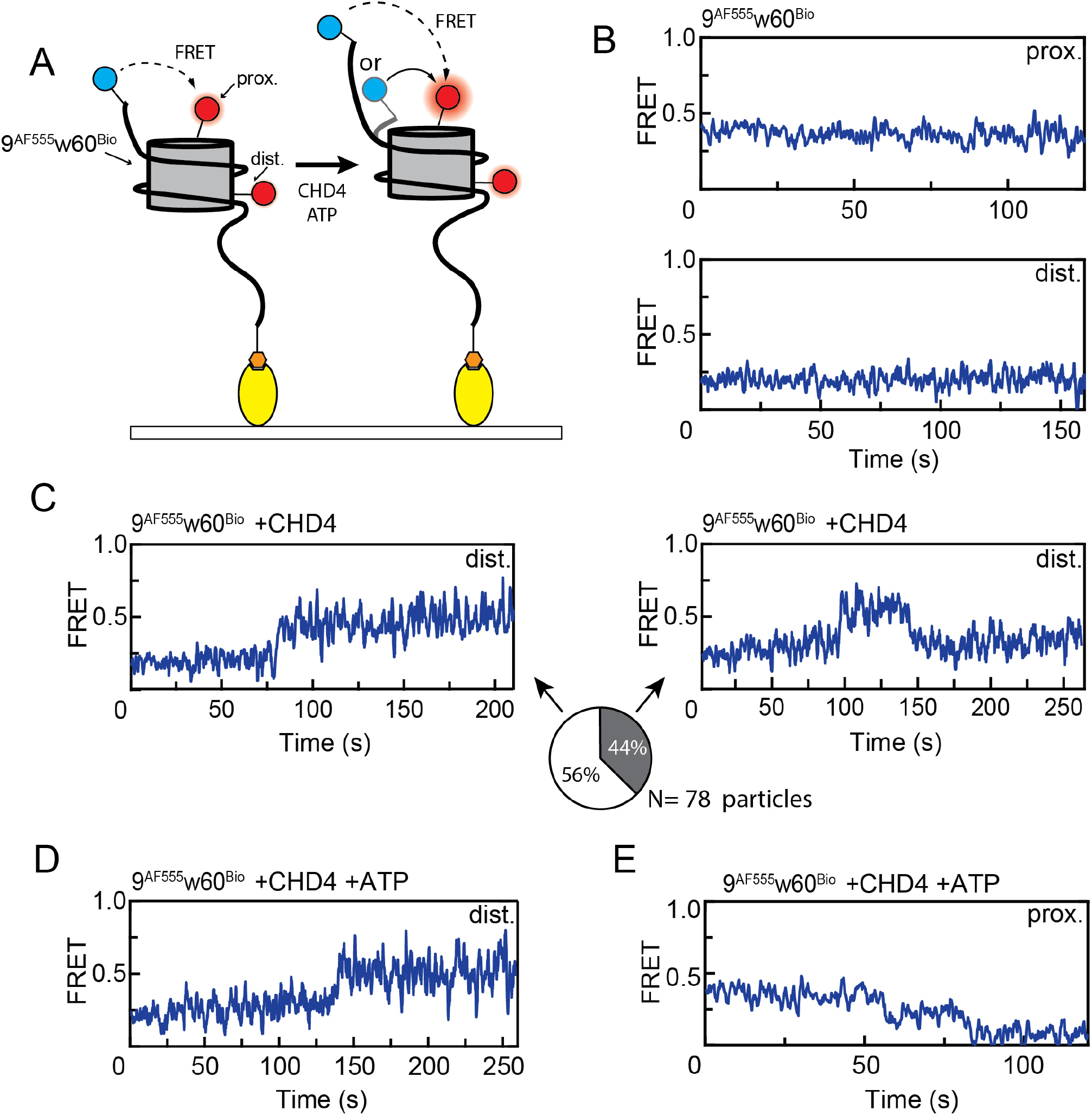
CHD4-induced similar FRET changes in extra-nucleosomal DNA at the TA-rich side. **A.** Schematic showing the labelling scheme for 9^AF555^w60^Bio^ and the two possible remodelling directions after treating with CHD4, depending on whether the ‘9’ end acts as an entry or an exit side. **B.** FRET vs time traces for 9^AF555^w60^Bio^ alone with a proximal (*top*) or distal (*bottom*) H2A AF647 label. **C.** FRET vs time trace for 9^AF555^w60^Bio^ bearing a distal AF647 label, showing an increase upon the addition of 2 nM CHD4. This newly established structure can be relatively stable (*left*) or can be transient, dropping back to the initial state (*right*). The proportions of each scenario (from 78 particles) are illustrated as a pie chart. **D**. FRET vs time trace for distally labelled 9^AF555^w60^Bio^ in presence of both 2 nM CHD4 and 1 mM ATP, showing an increase to a similar level to that of CHD4 alone. **E.** FRET vs time trace for proximally labelled 9^AF555^w60^Bio^, showing a stepwise drop upon treatment with 2 nM CHD4 and 1 mM ATP. In this case, the AF555 tag at the ‘9’ end is moving away from the histone octamer, in contrast to the direction of movement observed in D.

### The ‘9’ end of 9^AF555^w60^Bio^ can be stably remodelled as an exit but not an entry side

We also treated 9^AF555^w60^Bio^ nucleosomes with 2 nM CHD4 and 1 mM ATP. Surprisingly, we observed similar FRET changes as were observed with the addition of CHD4 alone; that is, increases in FRET from 0.2 to ~0.4, followed often by a return to 0.2, are observed for distally H2A-labelled particles (Figure 6D), although a small fraction of particles (15%) showed an increase to as high as ~0.6 (**Supplementary Figure 2D**). These results suggest that processive remodelling in which the ‘9’ end of the DNA acts as an entry site is not able to proceed, possibly because a 9-bp flanking sequence is too short to allow translocation without a destabilizing reduction in histone-DNA contacts. In contrast, 45% of the particles showed clear stepwise drops in FRET, indicating that 9^AF555^w60^Bio^ nucleosomes are indeed able to be remodelled – but in the opposite direction – with the ‘9’ end acting as an exit side (Figure 6E). Our gel-based assays using 9^AF555^w60 as a substrate support this interpretation: that this nucleosome can only be remodelled to a more symmetrical configuration (**Supplementary Figure 3**). Similarly, gel-based remodelling assays using the 0w30 and 30w0 substrates underscore the conclusion that CHD4 does not efficiently remodel shorter flanking DNA sequences towards the nucleosome (Figure 1E). The roughly equal (65:45) split between particles that display initial increases versus decreases in FRET is consistent with the idea that remodelling can occur in either direction in principle, most likely through CHD4 binding at each of its two possible binding sites.

Overall, these data demonstrate that nucleosome binding by CHD4 induces significant slow-timescale changes into extra-nucleosomal DNA, and that this process probably involves either twisting or sliding of the extra-nucleosomal DNA towards the histone octamer. The data also point towards a binding mode for CHD4 that is distinct from that of CHD1 (Sabantsev et al., 2019), in that there is not a significant displacement of several turns of DNA away from the nucleosome surface upon CHD4 binding.

## Discussion

### CHD4 is both similar to and distinct from CHD1

Prior to this work, little was known about the mechanism of CHD4-driven nucleosome remodelling. In contrast, a substantial body of work exists on yeast CHD1 structure and function. Yeast CHD1 and human CHD4 share 65% sequence similarity over the DNA translocase domain and of the 19 residues in the yCHD1 translocase domain that contact DNA, 12 are identical in hCHD4 (**Supplementary Figure 4**). Furthermore, structure prediction algorithms predict that the fold of the CHD4 translocase domain closely resembles that of yCHD1 (Yang et al., 2015).

EMSA data for CHD1 and CHD4 share similar features. Ryan *et al.* estimated the dissociation constant for a complex formed between CHD1 and a 0w47 nucleosome to be around 30 nM (Ryan et al., 2011), close to our observed value of 30–40 nM for CHD4. In addition, CHD1 also displays a second band shift at concentrations greater than 60 nM, which later was confirmed to represent a 2:1 complex in which a molecule of CHD1 is bound at each of the SHL2 sites (Farnung et al., 2017, Sundaramoorthy et al., 2018). The correspondence of our CHD4 gel-shift data with the yCHD1 data (Ryan et al., 2011) indicates that CHD4 probably also engages nucleosomes at the SHL2 and SHL– 2 sites.

Regulation of remodelling activity is likely to be distinct for the two enzymes, however. CHD1 has a C-terminal DNA-binding domain that binds to the other gyre of the nucleosome at SHL7, lifting two turns of DNA from the histone octamer surface (Farnung et al., 2017). This domain is absent from CHD4 and the deletion of the C-terminal third of CHD4 (which contains domains of unknown structure that mediate interactions with other proteins (Torrado et al., 2017, Marhold et al., 2004)) does not reduce its chromatin occupancy (Ramirez et al., 2012). In addition, we observed that CHD4 remodels symmetric substrates such as 30w30 more completely than asymmetric substrates (*e.g.*, 0w60), whereas CHD1 behaves in the opposite manner (McKnight et al., 2011, Stockdale et al., 2006). Interestingly, less CHD1 is required to remodel symmetric substrates if the DNA-binding domain is removed (McKnight et al., 2011), suggesting that this domain might act as a sensor of flanking DNA length that reduces the ability of CHD1 to move histone octamers to the end of a DNA sequence. Our data show that CHD4 does not possess this regulatory function, although the other subunits of the NuRD complex might well regulate CHD4 activity.

### CHD4-induced remodelling ejects 4–6 bp of DNA from the nucleosome in a concerted manner

We used smFRET to monitor distance changes between the DNA and the histone octamer during ATP-dependent nucleosome remodelling. When focusing on the exit end of the DNA, we observed FRET changes that represent translocation of 4–6 bp of DNA away from the histone octamer; these changes were consistent across a wide range of both CHD4 and ATP concentrations. Related observations have been made for the ISWI, RSC and CHD1 remodellers (Deindl et al., 2013, Harada et al., 2016, Qiu et al., 2017). In the case of ISWI, it was shown that each multiple-base-pair movement comprised a cluster of 1-bp steps that could be separated by lowering the ATP concentration. In contrast, however, a reduction in ATP concentration during CHD4 remodelling results only in an increase in the length of *t*_*pause*_ between the two steps; CHD1 has been reported to behave similarly (Qiu et al., 2017). These data indicate that the expulsion of 4–6 bp from the exit side is a concerted process.

Remodelling rate (defined as the length of *t*_*pause*_) was insensitive to CHD4 concentration, consistent with a processive mode of remodelling that does not involve dissociation/rebinding of CHD4. In contrast, the rate was highly dependent on ATP concentration, indicating that ATP turnover is a rate-limiting step for remodelling. We also observed that the distribution of *t*_*pause*_ times between exit-side translocation steps for CHD4 can be fitted to a gamma function and that the number of steps defined by the gamma function fit was essentially constant across a 2000-fold range of ATP concentrations. This distribution implies that the pause time comprises multiple rate limiting intermediate steps, and a fit of the distribution (Figure 4D) indicates that a reaction sequence of more than three steps best explains the data. That is, the data point to a model in which four or more ATP molecules are consumed during *t*_*pause*_, prior to the concerted exit-side translocation of 4–6 bp of DNA. Considering this inference together with our estimates of DNA translocation distance based on FRET changes, we propose that that each base pair movement might require ~1 ATP hydrolysis event. This value is in the range of mechano-chemical conversion efficiencies measured for other DNA helicases such as RecQ and PcrA, as well as for CHD1 (Rad and Kowalczykowski, 2012, Marians, 2000, Farnung et al., 2017).

### The binding of CHD4 induces dynamic conformational changes in nucleosomal DNA

In the process of making smFRET measurements at the DNA entry side, we unexpectedly observed that the binding of CHD4 alone was sufficient to trigger significant conformational changes in the flanking DNA. These changes were observed for both 9^AF555^w60^Bio^ and 0w(9^AF555^)60^Bio^ nucleosomes, indicating that the effect is independent of whether the flanking DNA is on the TA-rich or TA-poor side of the 601 sequence. Furthermore, these changes were gradual – a rise of ~0.2 FRET units over the course of 5–50 s that was followed in many cases by a more abrupt decrease to the original value. We interpret these changes as a reversible movement of flanking DNA towards the histone octamer, the energy for which is provided simply by binding of the remodeller. Following previously described models (Saha et al., 2006, Bowman, 2010), we propose that this movement occurs via a corkscrew-like twisting of the DNA – perhaps even of just one strand, to best maintain histone-DNA interactions. We note that no such changes were observed for 0^AF555^w60 nucleosomes in the presence of CHD4, consistent with the idea that the movement of the ‘0’ end into the nucleosome core would be too unfavourable because of the loss of histone-DNA interactions.

These data are consistent with the recent cryo-electron microscopy structure of Snf2 bound to the SHL2 site of a nucleosome in the absence of nucleotide. In this structure, the Snf2 induces a 1-bp translocation in one strand (the tracking strand, which orients 5’→3’ from SHL2 toward the dyad (Nodelman et al., 2017)) of the DNA at the SHL2 site to which the remodeller is bound. This effect appears to be propagated as far as the entry side (Li et al., 2019, Liu et al., 2017) and could very possibly extend into the flanking DNA. Analysis of histone-DNA crosslinking in CHD1-nucleosome complexes is also consistent with this idea with the observation by Winger *et al.* of a twist/shift of 1–3 bp at SHL5 upon binding of CHD1 at SHL2 (Winger et al., 2018). Indeed, distortions at the SHL2 site have been observed in structures of the nucleosome in isolation (Luger et al., 1999, McGinty and Tan, 2015, Muthurajan et al., 2003), suggesting that this location might be predisposed to be an initiation point for remodelling.

We observed comparable changes in entry-side FRET when incubating CHD4 and nucleosome in either the absence or presence of the non-hydrolysable nucleotide analogue AMP-PNP. This observation corroborates structural and smFRET data for Snf2 (Li et al., 2019) and also CHD1 crosslinking data (Winger et al., 2018). For both of these remodellers, the DNA distorting effect was stronger with apo or AMP-PNP-bound remodeller. In this case the two RecA-like lobes of the translocase domain are in a so-called ‘open’ conformation relative to each other (Liu et al., 2017, Hauk et al., 2010). In contrast, these changes in DNA structure are not observed for CHD1 or Snf2 in the presence of ADP⋅BeF_3_, an ATP analogue that induces a more ‘closed’ conformation involving a rotation of one lobe (Farnung et al., 2017, Civril et al., 2011, Li et al., 2019). Because distortion of the tracking strand of DNA appears to be a general feature of SF2-family helicases bound in the open/apo state to DNA (Singleton et al., 2007), and our AF555 fluorophores were attached on the extra-nucleosomal part of the tracking strand, our data suggest that CHD4 behaves in a similar manner as Snf2. That is, binding of CHD4 draws the tracking strand in from the entry side. An important aspect of the process that we report here is its slow timescale (up to tens of seconds). The reason for this timescale is not currently clear; it is possible that the required changes are coupled to slow, large-scale fluctuations of nucleosome structure that are gradually ‘captured’ by CHD4.

### A model for CHD4-mediated remodelling in which conformational changes at the entry and exit sides of the nucleosome are decoupled from each other

In the presence of both CHD4 and ATP, the FRET measured for 0w(9^AF555^)60^Bio^ increases from 0.2 to 0.8 or higher, indicating that DNA enters the nucleosome in a steady manner until the AF555 fluorophore prevents further translocation and the DNA is ‘reset’ to its starting position. This gradual increase contrasts sharply with the concerted ~5-bp translocations that we observe at the exit side and, taken together with the other available data, suggests a mechanism for CHD4 remodelling in which entry side and exit side movements are decoupled from each other (Figure 7).

**Figure 7.**
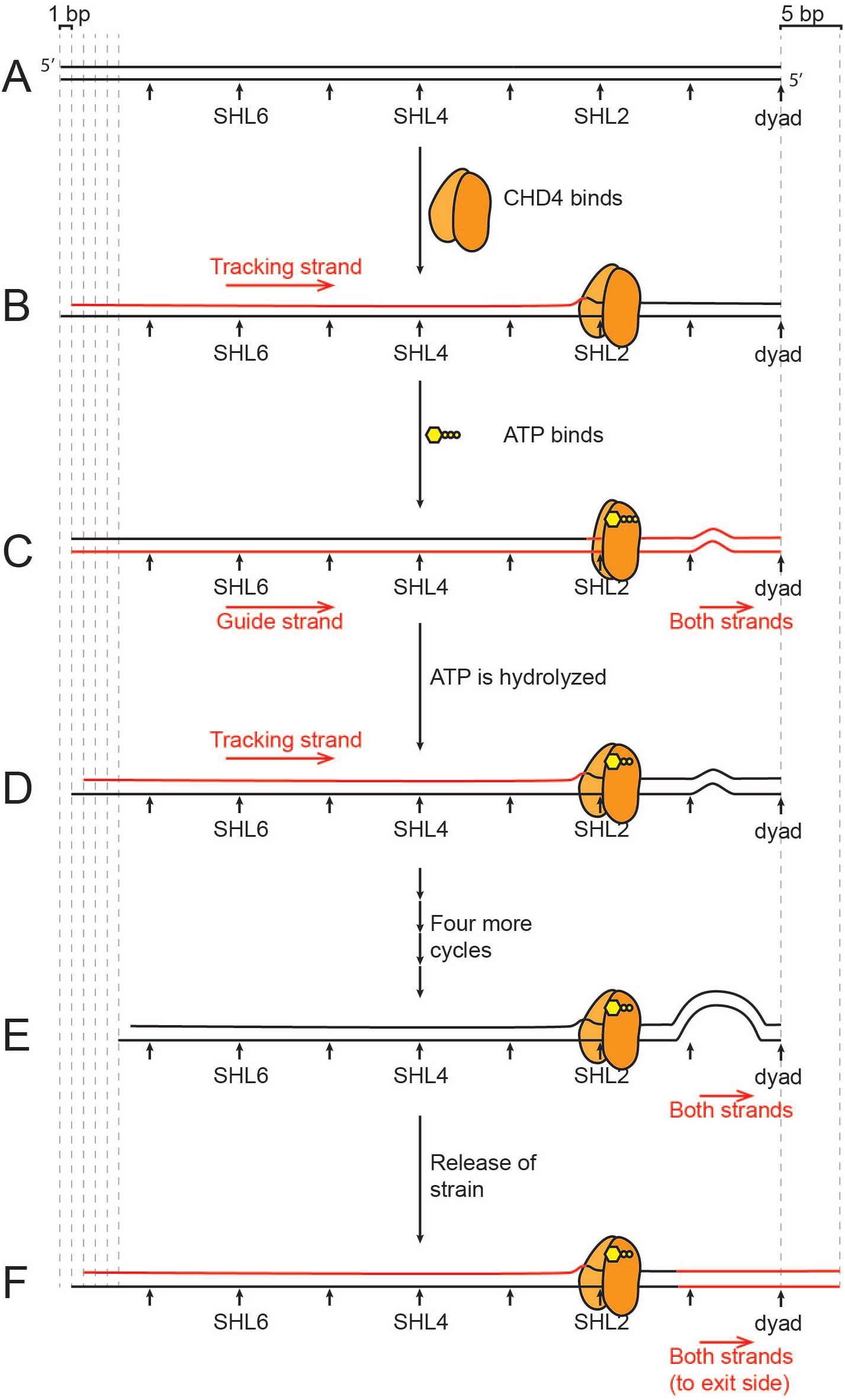
A model for CHD4-driven nucleosome sliding. **A.** Schematic of one half of a nucleosomal DNA sequence. SHL positions are indicated. **B.** Binding of the two-lobed ATPase domain of CHD4 induces a 1-bp shift in the so-called tracking strand of the DNA, creating a distortion that reaches from the SHL2 site all the way back to the 5’ end of the tracking strand. **C.** Binding of ATP induces a conformational change in CHD4, ‘closing’ the two lobes around the ATP. This change results in the guide strand ‘catching up’ to the tracking strand and then the movement of both strands by ~1 bp past CHD4, so that a small bulge (or other irregularity) forms in the region of the dyad. **D.** Hydrolysis of ATP drives a return of CHD4 to the open conformation, inducing a second 1-bp movement of the tracking strand, analogous to that in part B, and initiating a second cycle of the same process. **E.** Following four further cycles of ATP binding and hydrolysis, a large irregularity is built up near the dyad. **F.** The strain induced by this irregularity causes a concerted rearrangement of the DNA such that 5 bp are expelled from the exit side. This whole cycle can in principle be repeated many times.

We propose that, first, CHD4 binds SHL2 in an open conformation, inducing a 1-bp shift in the tracking strand that is propagated from SHL2 all the way back to the entry side (Figure 7B). This change is reversible, perhaps through CHD4 dissociation or through conformational changes in either CHD4 or the nucleosome. Second, binding of ATP closes the two lobes of CHD4, realigns the tracking and guide strands, and induces a distortion of the DNA between the SHL2 site and the dyad (Figure 7C). This process ‘relaxes’ the DNA between SHL2 and the entry side. Third, following ATP hydrolysis, the two lobes of CHD4 return to the open position, establishing a new 1-bp translocation of the tracking strand that pulls a nucleotide from the entry side (Figure 7D). This is consistent with the available structural and biochemical data that show an open position for SF2 family remodellers in both the apo and ADP bound states (Li et al., 2019). This cycle continues until a ~5-bp distortion is built up near the dyad (Figure 7E). The nature of this strained state is not yet known, but it could either be a small loop or a small segment of overwound DNA helix (Cairns, 2007). Eventually, the strain induced by this distortion becomes too great to be maintained and is released at the exit side by a concerted twist and translocation of the DNA (Figure 7F). This entire process is repeated in a processive mechanism that moves DNA relative to the histone octamer.

In conclusion, we have carried out the first mechanistic analysis of CHD4-driven chromatin remodelling and our data lead to a model in which processive CHD4 action builds up strain in the DNA through 4–6 successive ATP-dependent translocations of 1 bp, and the strain is released by a concerted expulsion of those base pairs from the exit side. This model synthesizes a wide range of recent structural and mechanistic measurements on several other chromatin remodelling enzymes, suggesting common mechanistic features across SF2-family remodellers, while at the same time indicating that some differences exist in the precise mechanisms of action of each remodeller.

## Materials and methods

### Histone purification and labelling

Recombinant human histones were expressed in Escherichia coli (*E. coli*) BL21 (DE3) pLysS cells and purified from inclusion bodies. Competent cells were transformed with histone expression plasmids (pET28a, Novagen) and grown in LB medium at 37 °C until reaching an OD600 of 0.6. Protein expression was induced by the addition of 1 mM IPTG for 4 h at 37 °C. Cell pellets were harvested and lysed by sonication in histone lysis buffer containing 50 mM Tris-HCl pH 7.5, 100 mM NaCl, 1 mM beta-mercaptoethanol (BME), and the insoluble fraction after clarification (30 min at 15, 000 ×g at room temperature) was washed twice with histone lysis buffer containing 1% (v/v) Triton X-100, and then twice without Triton X-100. The pellet was dissolved in 10 mL of unfolding buffer (20 mM Tris-HCl pH 7.5, 6 M guanidinium HCl and 1 mM dithiothreitol (DTT)) per L of culture by stirring at room temperature overnight. Resuspended pellets were then centrifuged at 20,000 ×g for 30 min at 4 °C. Filtered supernatants were injected onto a preparative Vydac protein and peptide C18 column (300 Å pore size, Catalogue No. 218TP1022) at a flow rate of 7 mL/min (20–70% acetonitrile over 40 min, 0.1% TFA). The fractions containing the target protein (as judged by UPLC-MS analysis) were freeze-dried and stored at –20 °C in sealed containers.

To generate H2A site-specifically labelled with AF647, a T120C mutant was generated by site-directed mutagenesis. This construct was purified by reverse-phase HPLC using the same method employed for the wild-type proteins. After lyophilization, 200 nmole of H2A T120C was dissolved in 0.5 mL of unfolding buffer containing 1 mM tris(2-carboxyethyl)phosphine (TCEP), followed by degassing. The solution was then mixed directly with a 5× molar excess of Alexa Fluor™ 647 C_2_ maleimide (Thermo Fisher Scientific A20347), and then incubated at room temperature for 10 min and then at 4 °C overnight. The reactions were quenched via the addition of 30 mM BME and then purified via gel filtration on a Superdex 200 10/300 column in 20 mM Tris, pH 7.0, 7 M guanidine HCl, 0.1% (v/v) BME. Purified labelled H2A was dialyzed against deionized water with 0.05% (v/v) BME overnight at 4 °C and lyophilized for long term storage. Labelling efficiency was ~50%, determined using method described in (Kim et al., 2008).

### Nucleosomal DNA preparation

DNA oligonucleotides were made by PCR from a plasmid containing a 601 positioning sequence (Lowary and Widom, 1998). The PCR primers contained 5’ AF555 or biotin-TEG modifications (Integrated DNA Technologies, Singapore) to install these modifications at the indicated locations. The PCR products were first concentrated by ethanol precipitation and were redissolved in 1× TE buffer (10 mM Tris-HCl pH 8, 1 mM EDTA), and then cleaned up using phenol:chloroform:isoamyl alcohol (25:24:1) extraction, followed by a chloroform wash to remove residue phenol. After isopropanol precipitation and washing with 70% (v/v) ethanol, the DNA pellet was dissolved in 1× TE and separated on a 0.5× TBE 5% polyacrylamide gel. The DNA band with the desired size was cut from the gel and electroeluted into 0.5× TBS at room temperature. The final DNA product was concentrated by ethanol precipitation overnight and the resulting pellet was dissolved in 1× TE.

### Nucleosome preparation

Histone octamer was reconstituted as described in (Luger et al., 1999). Mononucleosomes were assembled by salt gradient dialysis using a double dialysis method (Thastrom et al., 1999). After mixing labelled octamer and DNA at a 1:0.95 molar ratio in 10 mM Tris-HCl pH 7.5, 2 M NaCl, 1 mM EDTA, the mixture was loaded into a small dialysis button, which was then placed into a dialysis bag containing 30 mL of 10 mM Tri-HCl pH 7.5, 2 M NaCl, 1 mM EDTA and 0.1 mM DTT. The bag was then dialysed against 2 L of 1× TE containing 0.1 mM DTT overnight at room temperature.

The next day, the dialysis button was dialyzed further against 10 mM Tris pH 7.5, 2.5 mM NaCl and 0.1 mM DTT. Content in the dialysis button was harvested and the nucleosome quality was checked by 0.5× TBE 5% polyacrylamide gel electrophoresis at 150 V in the cold room.

### Expression and purification of FLAG-CHD4 in HEK293 cells

Suspension-adapted HEK Expi293F™ cells (Thermo Fisher Scientific, Waltham, MA, USA) were grown to a density of 2× 10^6^ cells/mL in Expi293™ Expression Medium (Thermo Fisher Scientific). pcDNA3.1 plasmids encoding for FLAG-CHD4 were transfected into cells using linear 25-kDa polyethylenimine (PEI, Polysciences, Warrington, PA, USA). 50 µg of the DNA mixture was first diluted in 2.7 mL of PBS and vortexed briefly. 100 µg of PEI was then added, and the mixture was vortexed again, incubated for 20 min at room temperature, and then added to 25 mL of HEK cell culture. The cells were incubated for 65 h at 37 °C with 5% CO_2_ and horizontal orbital shaking at 130 rpm.

Cells were harvested and washed twice with PBS, centrifuged (300 g, 5 min), snap-frozen in liquid nitrogen and stored at –80 °C. Lysates were prepared by sonicating thawed cell pellets in 0.5 mL of lysis buffer [50 mM Tris/HCl, 500 mM NaCl, 1% (v/v) Triton X-100, 1× cOmplete EDTA-free protease inhibitor (Roche, Basel, Switzerland), 100 mM ATP, 0.2 mM DTT, pH 7.9], and then clarifying the lysate via centrifugation (≥ 16 000 g, 20 min, 4 °C). The cleared supernatant was used for FLAG-affinity pulldowns.

200 µL of anti-FLAG Sepharose 4B beads [Biotool, Houston, TX, USA; pre-equilibrated with 50 mM Tris-HCl, 150 mM NaCl, 0.1% (v/ v) Triton X-100, 1× cOmplete EDTA-free protease inhibitor, 0.2 mM DTT, pH 7.5] was added to 15 mL of cleared HEK cell lysate. The mixtures were incubated overnight at 4 °C with orbital rotation. Post-incubation, the beads were washed with 5× wash buffer [50 mM Tris-HCl, 150 mM NaCl, 0.5% (v/v) IGEPAL CA630, 100 mM ATP, 0.2 mM DTT, pH 7.5]. Bound proteins were eluted by 5× 400 µL treatment with ‘elution’ buffer (10 mM HEPES, 150 mM NaCl, 150 µg/mL 3× FLAG peptide (MDYKDHDGDYKDHDIDYKDDDDK), pH 7.5) for 1 h at 4 °C. Concentrations of CHD4 in the elution fractions were estimated using densitometry (ImageJ) by loading onto a SDS-PAGE and staining with SYPRO® Ruby.

### NuRD purification

NuRD was purified as described in (Low et al., 2016). Briefly, GST-FOG1(1–45) was overexpressed in *E. coli* BL21(DE3) cells. The cells were lysed via sonication in GST binding buffer (50 mM Tris, 150 mM NaCl, 0.1% BME, 0.5 mM PMSF, 0.1 mg/ml lysozyme, 10 µg/ml DNase I, pH 7.5) and clarified via centrifugation (≥16,000× g, 20 min, 4 °C). The cleared supernatant was then incubated with pre-equilibrated glutathione-Sepharose 4B beads (GE Healthcare) for 1 h at 4 °C. The beads were then washed in GST wash buffer (20 CV, 50 mM Tris, 500 mM NaCl, 1% (v/v) Triton X-100, 1mM DTT, pH 7.5) and then NuRD binding buffer (10 CV, 50 mM HEPES-KOH, 150 mM NaCl, 1% v/v Triton X-100, 1 mM DTT, 1× cOmplete^®^ protease inhibitor (Roche), pH 7.4). The beads were then used for NuRD pulldown experiments.

MEL cell nuclear extracts were prepared by incubating the thawed cell pellets with hypotonic lysis buffer (5 ml/g of cells; 10 mM HEPES-KOH, 1.5 mM MgCl_2_, 10mM KCl, 1 mM DTT, cOmplete^®^ protease inhibitor, pH 7.9) for 20 min at 4 °C. IGEPAL^®^ CA-630 was then added (final concentration, 0.6% v/v), and the cells were then further incubated for 10 min. The mixture was then vortexed for 10 s and then centrifuged (3,300× g, 5 min). The cytoplasmic supernatant was discarded, and the nuclear pellet was gently washed once with lysis buffer (+ 0.6% (v/v) IGEPAL^®^ CA-630). The washed nuclear pellet was resuspended in NuRD binding buffer (3 mL/g of cells), then lysed by sonication, and incubated on ice for 30 min to allow the chromatin to precipitate. The nuclear extract was then clarified via centrifugation (≥16,000× g, 20 min, 4 °C), and the cleared supernatant was incubated with Streptavidin beads (a preclearing step for FOG1(1–45) peptide affinity purification) before incubating with the above FOG1 affinity resins overnight at 4 °C. Post-incubation, the nuclear extract was then washed with 20 CV of NuRD wash buffer 1 (50 mM HEPES-KOH, 500 mM NaCl, 1% (v/v) Triton X-100, 1 mM DTT, pH 7.4) and then 10 CV of

NuRD wash buffer 2 (50mM HEPES-KOH, 150mM NaCl, 0.1%(v/v) Triton X-100, 1mM DTT, pH7.4). Captured proteins were eluted with GST-FOG1(1–45) elution buffer (50 mM reduced glutathione, 50 mM HEPES-KOH, 150 mM NaCl, 0.1% Triton X-100, 1mM DTT, pH 8.0) for 30 min at 4 °C. This elution step was repeated at least twice to ensure complete elution.

### Nucleosome repositioning assay

Reactions contained 50 nM of either labelled or non-labelled nucleosomes, 50 mM Tris pH 7.5, 50 mM NaCl, 3 mM MgCl_2_, and the enzyme concentrations varied from 0 to 10 nM. The reactions were incubated at 37 °C and then stopped by placing them on ice and the addition of 0.5 µg salmon sperm DNA and 4% (w/v) sucrose prior to electrophoresis on 0.5× TBE 5% polyacrylamide gels. Gels were stained with 1× SYPRO® Gold and then imaged by on an FLA-9000 laser scanner.

### Nucleosome pull-down assay

HEK Expi293F™culture expressing FLAG-CHD4 (3 mL) was prepared as described above. The cell pellet was then lysed in the same way and loaded onto anti-FLAG beads. After five washes, the beads were washed three times again with 10 mM Tris pH 7.5, 2.5 mM NaCl and 0.1 mM DTT, and split into 3 aliquots, and each was incubated with the same amount of nucleosome (~3 pmol in 10 mM Tris pH 7.5, 2.5 mM NaCl and 0.1 mM DTT) at 4 °C overnight. The next day, the proteins were eluted using the method as above and equivalent amounts of input and elution were checked by SDS-PAGE.

### EMSA of nucleosome-CHD4 interaction

Each reaction contained 60 nM of AF647 labelled nucleosomes, 50 mM Tris pH 7.5, 50 mM NaCl, 3 mM MgCl_2_, and the enzyme concentrations as indicated in the figures. The reactions were incubated on ice for 60 min, protected from light, and then mixed with 4% (w/v) sucrose prior to electrophoresis on 0.5× TBE 5% polyacrylamide gels. Gels were then imaged on an FLA-9000 laser scanner.

### Single-molecule instrument setup

An Olympus IX-71 based model was modified to build an objective type total internal reflection fluorescence (TIRF) microscope to record single-molecule movies. A coherent Sapphire green (532 nm) laser was used to excite donor (AF555) molecules by focusing onto a 100× oil immersed objective and scattered light was removed using a 560-nm long pass filter. Donor and acceptor (AF647) signals were collected at 565 and 665 nm using a band pass filter (560–600) nm and a long pass filter at 650 nm, respectively. Then, both signals were first split by a 638-nm dichroic mirror using Photometrics Dual View (DV-2) and then were focused onto a CCD camera (Hamamatsu C9 100-13), simultaneously. Single-molecule movies were recorded at 5 frames per second.

### Preparation of PEG-ylated coverslips

First, quartz coverslips were sonicated with 2–5 M KOH for 20 min and rinsed with double distilled water (ddH_2_O). Second, aminosilation of coverslips were carried out in a mixture of 100 mL water and 1% (v/v) aminopropylsilane (Alfa Aesar, A10668, UK). Third, PEGylation was carried out by incubating a mixture of biotinPEG-SVA and mPEG-SVA (Laysan Bio, AL, USA) in the ratio of 1:20 prepared in 50 mM MOPS at pH 7.5 on the top of the silanized coverslip for 3-4 h. Finally, PEG-ylated coverslips were rinsed with ddH_2_O, dried with dry nitrogen and stored under dry nitrogen gas at –20 °C.

### Single-molecule FRET experiments

Immuno-pure neutravidin solution was prepared in imaging buffer (40 mM Tris-HCl, pH 7.5, 12 mM HEPES, pH 7.5, 3 mM MgCl_2_ and 60 mM KCl, 0.32 mM EDTA, 10% (v/v) glycerol, 0.02% (v/v) IGEPAL (Sigma Aldrich) and spread on the top of dry PEG-ylated coverslip for 10 min. Then, polydimethylsiloxane (PDMS) was sandwiched on the top of the neutravidin coated coverslip to create a microscopic channel. Then, blocking buffer (prepared by mixing 1% (v/v) Tween-20 in imaging buffer) was injected onto the microscopic channel, in order to reduce non-specific binding of proteins on the surface, and incubated for 10-15 min.

Different mononucleosome samples labelled with FRET pair fluorophores and biotin were diluted to 50 pM in imaging buffer and injected into the flow chamber using a syringe pump (ProSense B.V.); the mixture was then incubated for 5–10 min. Unbound sample was removed by flowing imaging buffer through the chamber. Next, an oxygen-scavenging system (OSS) consisting of protocatechuic acid (PCA, 2.5 mM) and protocatechuate-3,4-dioxygenase (PCD, 50 nM) in imaging buffer were flowed across the surface to reduce photobleaching of the fluorophores. Trolox (2 mM) was also added to reduce photoblinking of dyes. Multiple movies were then recorded to measure the distribution of free nucleosomes. For CHD4 binding assays, a mixture of CHD4 (2–20 nM) diluted in imaging buffer (containing OSS) was injected for 10–20 s while a movie was recorded continuously. For remodeling assays, additional ATP (0.01–20 mM) was included in the imaging buffer. The CHD4 binding or remodeling mixture reached the reaction chamber in 10–20 s and a movie was recorded continuously at room temperature (20 ± 1 °C) for 3–5 min (until acceptor dyes photobleached).

### Data analysis

Single-molecule intensity time trajectories were generated in interactive data language (IDL) and these trajectories were analyzed in MATLAB using home written scripts (https://cplc.illinois.edu/software/). An approximate FRET value is measured as the ratio of acceptor intensity to the sum of the donor and acceptor intensities after correcting for cross-talk between donor and acceptor channels.

Since the acceptor dye was on the H2A subunit of the histone octamer, it gives rise to heterogeneity in FRET population, because three different labelling states exist: i) acceptor dye proximal to donor dye, yielding a mid-FRET state; ii) acceptor dye distal to donor dye, giving a low-FRET state; iii) two acceptor dyes on both H2A subunits, yielding a high-FRET state. For example, we observed a distribution that could be modelled by three Gaussians with mean FRET values of 0.78, 0.67 and 0.42for exit side 0w60 nucleosomes. Similarly, for entry side labelled nucleosome (0W9AF555-60) we observed peaks with means of 0.62, 0.38 and 0.23. In this study, we focused on the molecules in the mid-FRET state. In the FRET trace analysis, we chose molecules showing a mean FRET value in the range of 0.55 to 0.7 and displaying a single acceptor photobleaching step. In some cases, we also selected populations showing mean FRET values in the range of 0.35 to 0.5 and showing a single step acceptor photobleaching, in order to analyze distal side dynamics. These selection criteria allowed us to distinguish between nucleosomes bearing one H2A labelled subunit. Similar criteria with different FRET cut-off values were applied to selectmid-FRET traces in different nucleosomes(n = 3, 6 and 9) for calibration purpose. For entry site labelled nucleosomes, distally labelled mono-nucleosomes were chosen for statistical analysis.

To measure the rate of remodeling reaction at different ATP or CHD4 concentrations, histograms were generated in each condition and fit with a Gaussian, where the peak of the curve fit was taken as the mean time for remodeling by CHD4.

### Gamma distribution

To measure the number of steps involved in a remodelling reaction, a Gamma distribution was applied to time-binned histograms of the form:

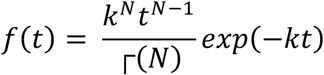

Where *k* is the rate of the reaction and *N* is the number of steps hidden in the reaction.

## Acknowledgements

The work was funded by the following grants from the National Health and Medical Research Council of Australia: APP1012161, APP1063301, APP1126357 and a fellowship from the same organization to JPM (APP1058916). AMvO is an Australian Research Council Laureate Fellow.

## Author Contributions

Conceptualization, Y.Z., B.P.P., A.M.O. and J.P.M.; Methodology, Y.Z., B.P.P., D.P.R., J. K.K.L., C.F., K.P. and M.J.B.; Software, B.P.P.; Investigation Y.Z., B.P.P.; Resources, R.J.P., A.M.O. and J.P.M.; Writing – Original Draft, Y.Z. and J.P.M.; Writing – Reviewing and Editing, Y.Z., B.P.P., M.J.B., R.J.P., A.M.O. and J.P.M.; Supervision, A.M.O. and J.P.M.

