## Supplementary Figures for "CHD4 slides nucleosomes by decoupling entry- and exit-side DNA translocation"

**Supplementary Information**

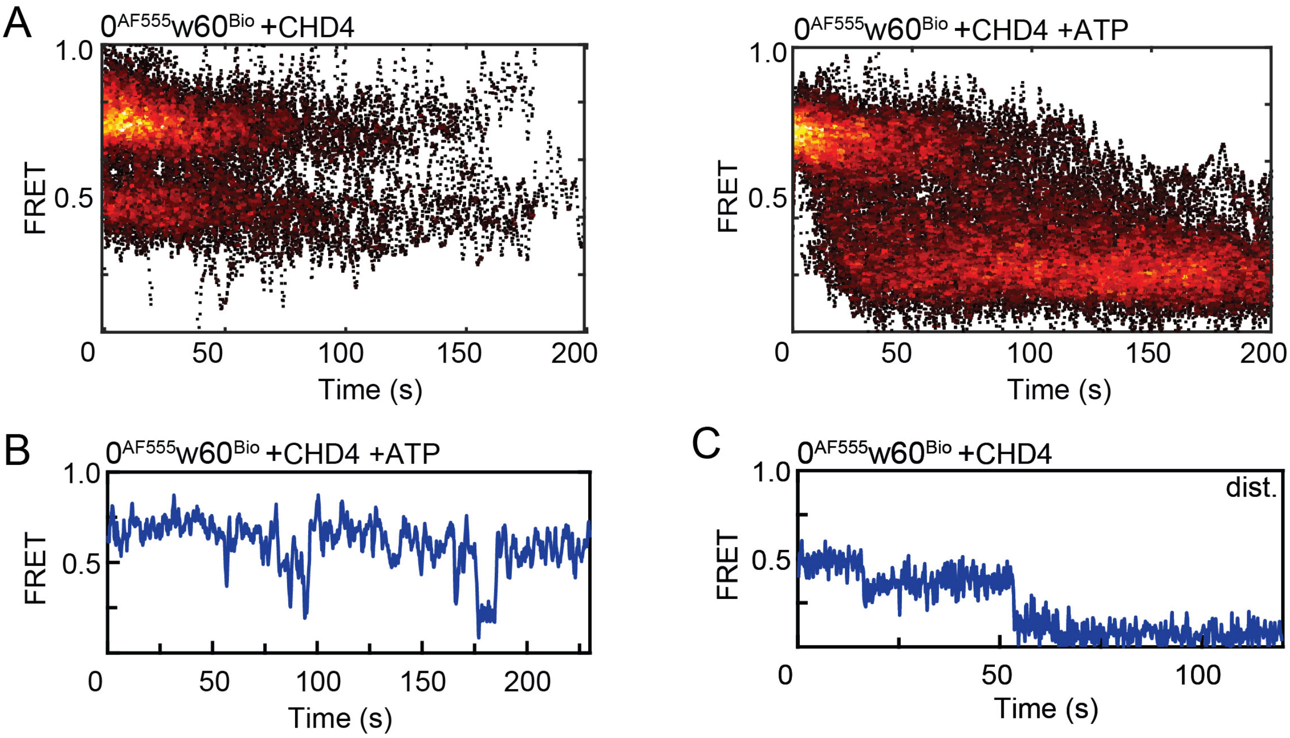


**Figure S1. Control smFRET experiments for CHD4-driven remodelling of $0^{AF555}w60^{Bio}$ nucleosome. A.** Heatmap of both proximally and distally labelled $0^{AF555}w60^{Bio}$ nucleosomes showing the time dependence of FRET upon incubation with either CHD4 alone (*left panel*, N = 105) or with 2 nM CHD4 in the presence of 1 mM ATP (*right panel*, N = 79). **B.** FRET vs time trace for $0^{AF555}w60^{Bio}$, showing the transient fluctuations that were observed occasionally in the presence of both CHD4 and ATP. **C.** FRET vs time trace for a distally labelled $0^{AF555}w60^{Bio}$ nucleosome, showing a step-wise decrease when incubated with 1 mM ATP and 2 nM CHD4.

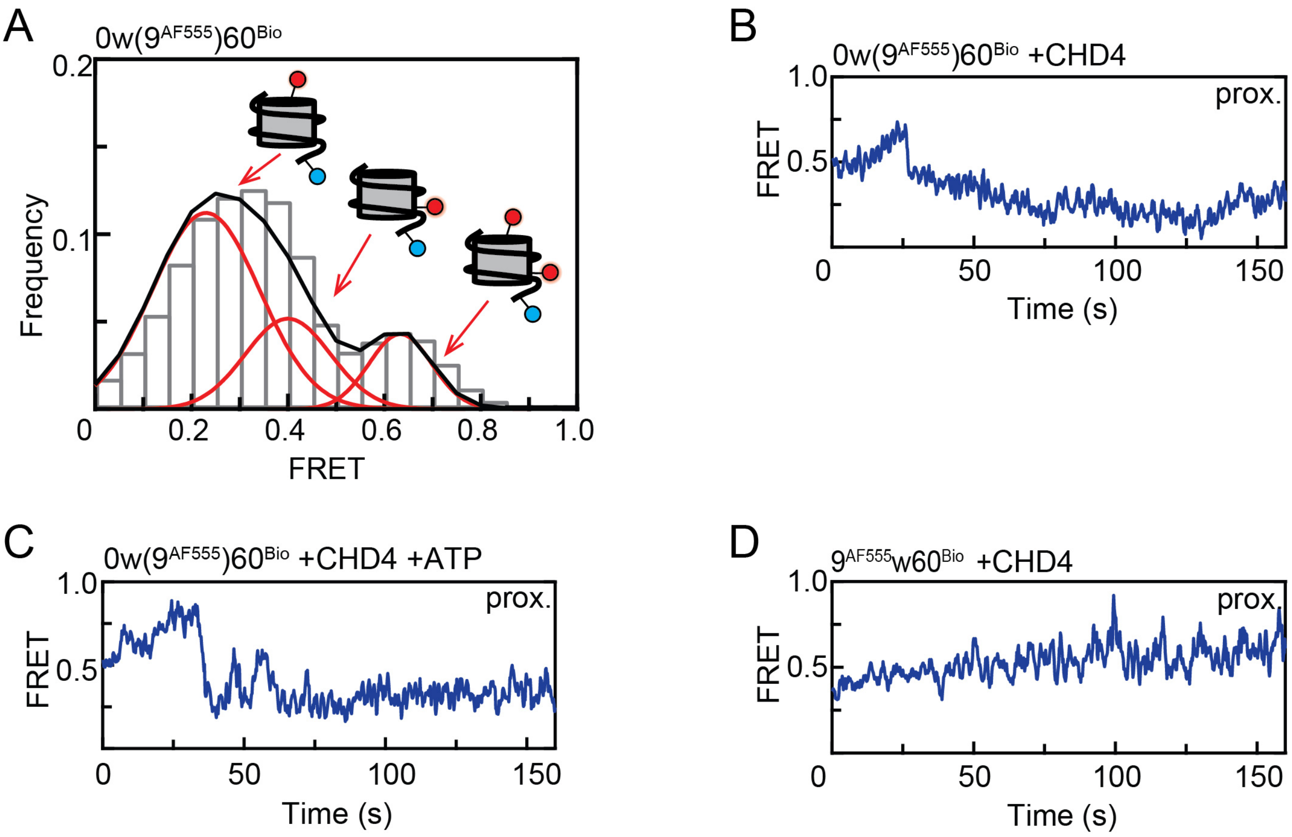


**Figure S2. Pre-reaction distribution of nucleosomal FRET states for 0w(9$^{AF555}$)60$^{Bio}$ nucleosomes and FRET traces of proximally labelled 0w(9$^{AF555}$)60$^{Bio}$ and 9$^{AF555}$w60$^{Bio}$ nucleosomes undergoing CHD4 binding or remodelling. A.** FRET distribution for 0w(9$^{AF555}$)60$^{Bio}$ nucleosomes (N = 97). The peak FRET values 0.2, 0.4, and 0.65 (obtained from a Gaussian fit, black line) correspond to particles bearing a AF647 tag at proximal, distal or both H2A subunits. **B**. FRET trace of 0w(9$^{AF555}$)60$^{Bio}$ bearing a proximal AF647 label, showing a gradual increase and then decrease upon binding of CHD4 (2 nM). The high FRET state is transient and quickly drops back to 0.4 and then subsequently 0.2. **C.** FRET vs time trace for 0w(9$^{AF555}$)60$^{Bio}$ bearing a proximal AF647 label, in the presence of both CHD4 (2 nM) and ATP (10 μM). **D.** FRET vs time trace for 9$^{AF555}$w60$^{Bio}$ bearing a proximal AF647 label, in the presence of CHD4 (2 nM). A gradual increase is observed over time.

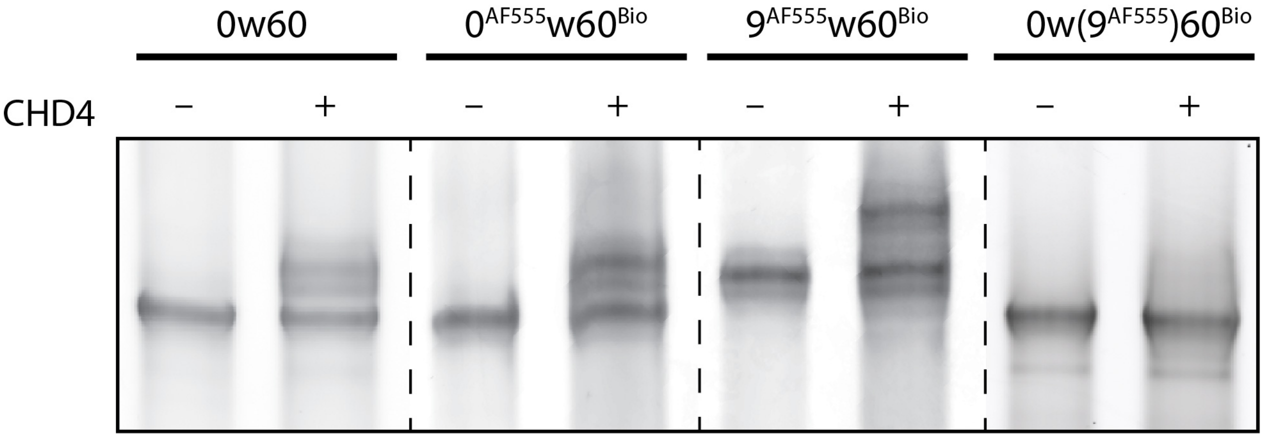


**Figure S3. Gel-based nucleosome repositioning assays for the nucleosomes used in smFRET assays.** Nucleosomes containing both AF555 on DNA and AF647 on H2A were remodelled by 10 nM CHD4 in the presence of 1 mM ATP. A positive control of unlabelled 0w60 was included.

```
CHD4_HUMAN LHPYQMEGLNWLRFSWAQGTDTILADEMGLGKTVQTAVFLYSLYKEGHSKGPFLVSAPLS 785
CHD1_YEAST LRDFQLTGINWMAFLWSKGDNGILADEMGLGKTVQTVAFISWLIFARRQNGPHIIVVPLS 435
CHD1_HUMAN LRDYQLNGLNWLAHSWCKGNSCILADEMGLGKTIQTISFLNYLFHEHQLYGPFLLVVPLS 540

CHD4_HUMAN TIINWEREFEMWAPDMYVVTYVGDKDSRAIIRENEFSFEDNAIRGGKKASRMKKEASVKF 845
CHD1_YEAST TMPAWLDTFEKWAPDLNCICYMGNQKSRDTIREYEFYTNPR----------AKGKKTMKF 485
CHD1_HUMAN TLTSWQREIQTWASQMNAVVYLGDINSRNMIRTHEWTHH--------------QTKRLKF 586

CHD4_HUMAN HVLLTSYELITIDMAILGSIDWACLIVDEAHRLKNNQSKFFRVLNGYSLQHKLLLTGTPL 905
CHD1_YEAST NVLLTTYEYILKDRAELGSIKWQFMAVDEAHRLKNAESSLYESLNSFKVANRMLITGTPL 545
CHD1_HUMAN NILLTTYEILLKDKAFLGGLNWAFIGVDEAHRLKNDDSLLYKTLIDFKSNHRLLITGTPL 646

CHD4_HUMAN QNNLEELFHLLNFLTPERFHNLEGFLEEFADIAKEDQIKKLHDMLGPHMLRRLKADVFKN 965
CHD1_YEAST QNNIKELAALVNFLMPGRFTIDQEIDFENQDEEQEEYIHDLHRRIQPFILRRLKKDVEKS 605
CHD1_HUMAN QNSLKELWSLLHFIMPEKFSSWEDFEEEHGKG-REYGYASLHKELEPFLLRRVKKDVEKS 705

CHD4_HUMAN MPSKTELIVRVELSPMQKKYYKYILTRNFEALNARGGGNQVSLLNVVMDLKKCCNHPYLF 1025
CHD1_YEAST LPSKTERILRVELSDVQTEYYKNILTKNYSALTAGAKGGHFSLLNIMNELKKASNHPYLF 665
CHD1_HUMAN LPAKVEQILRMEMSALQKQYYKWILTRNYKALSKGSKGSTSGFLNIMMELKKCCNHCYLI 765

CHD4_HUMAN PVAAMEAP-KMPNGMY----DGSALIRASGKLLLLQKMLKNLKEGGHRVLIFSQMTKMLD 1080
CHD1_YEAST DNAEERVLQKFGDGKMTRENVLRGLIMSSGKMVLLDQLLTRLKKDGHRVLIFSQMVRMLD 725
CHD1_HUMAN KPPDNNEF-------YNKQEALQHLIRSSGKLILLDKLLIRLRERGNRVLIFSQMVRMLD 818

CHD4_HUMAN LLEDFLEHEGYKYERIDGGITGNMRQEAIDRFNAPGAQQFCFLLSTRAGGLGINLATADT 1140
CHD1_YEAST ILGDYLSIKGINFQRLDGTVPSAQRRISIDHFNSPDSNDFVFLLSTRAGGLGINLMTADT 785
CHD1_HUMAN ILAEYLKYRQFPFQRLDGSIKGELRKQALDHFNAEGSEDFCFLLSTRAGGLGINLASADT 878

CHD4_HUMAN VIIYDSDWNPHNDIQAFSRAHRIGQNKKVMIYRFVTRASVEERITQVAKKKMMLTHLVVR 1200
CHD1_YEAST VVIFDSDWNPQADLQAMARAHRIGQKNHVMVYRLVSKDTVEEEVLERARKKMILEYAIIS 845
CHD1_HUMAN VVIFDSDWNPQNDLQAQARAHRIGQKKQVNIYRLVTKGSVEEDILERAKKKMVLDHLVIQ 938
```

**Figure S4. Sequence alignment of the ATPase domain of human CHD4 with human and yeast CHD1.** Of the 19 residues that make contact with the DNA in the structure of CHD1 bound to the nucleosome (PDB: 5O9G, (Farnung et al., 2017), marked in *yellow*), conserved residues are highlighted by *black boxes*. Additional conservation between human CHD1 and CHD4 is highlighted by *red boxes*.
